# Sample barcoding-associated technical variation in probe-based single-cell RNA sequencing

**DOI:** 10.64898/2026.04.06.716804

**Authors:** Jackson A. Weir, Yonit Krebs, Fei Chen

**Affiliations:** Broad Institute of MIT and Harvard, Cambridge, MA, USA; Biological and Biomedical Sciences Program, Harvard University, Cambridge, MA, USA; Department of Molecular and Cellular Biology, Harvard University, Cambridge, MA, USA; Department of Stem Cell and Regenerative Biology, Harvard University, Cambridge, MA, USA

## Abstract

Probe-based single cell RNA sequencing approaches are increasingly becoming a technology of choice for profiling gene expression at scale and in archival tissues. The 10x Genomics Flex v1 assay enables cost-effective and high-sensitivity single-cell RNA sequencing by splitting samples across up to 16 uniquely barcoded probe sets before pooling and loading onto a single lane of a microfluidic chip. A natural consequence of this design is to leverage probe set barcoding as a sample barcoding strategy for case-control experiments. However, we observed that Flex v1 probe set barcode identity drives substantial technical variation between probe set barcodes, an effect that is reproducible across lanes and independent datasets. When Flex v1 probe set barcodes are confounded with biological sample identity, a concerning number of differentially expressed genes at standard thresholds are false positives. The Flex v2 assay, which decouples sample barcoding from probe set hybridization, significantly reduces this artifact. As the field continues to expand adoption of probe-based assays, our findings introduce probe set barcoding as an underappreciated source of technical variation in single-cell assays and emphasize the importance of experimental design when using probe-based sequencing technologies.

## Main text

Single-cell RNA sequencing has become a central tool for measuring cellular phenotypes across disciplines of biology. Increasingly, probe-based single cell profiling, most notably the Flex Gene Expression assay from 10x Genomics, has offered the promise of higher transcript measurement sensitivity with lower per cell costs^1,2^. These technological advances have motivated enormous data collection efforts^3,4^, have enabled exciting technology development^5^, and have produced a technology of choice for generating training data for virtual cell models^6,7^.

The scalability driving the broad appeal of Flex is enabled by probe set-level barcoding. Flex uses established transcriptome sets to generate pairs of oligonucleotide probes that can hybridize to neighboring regions of a gene transcript and ligated together into a single DNA fragment for further amplification and sequencing. In Flex v1, the transcriptome probe kit is available in configurations supporting up to 16 uniquely barcoded probe sets, each comprising probe pairs that cover the full transcriptome but carrying distinct 8 bp probe barcode sequences. Samples can be hybridized with probe sets carrying different probe barcodes, and subsequently pooled and co-loaded onto the microfluidic chip (Figure 1A). Because each ligated probe carries a probe barcode, droplets that contain multiple cells from hybridization reactions with uniquely barcoded probe sets can be computationally demultiplexed into individual cells. This enables superloading of the microfluidic chip and cheaper per cell processing costs. Naturally, this design encourages the use of probe set barcodes as biological sample barcodes to process many samples on a single lane of the microfluidic chip. For workflows aiming to make case-control comparisons across uniquely barcoded probe sets, batch effect should be minimal.

**Figure 1.**
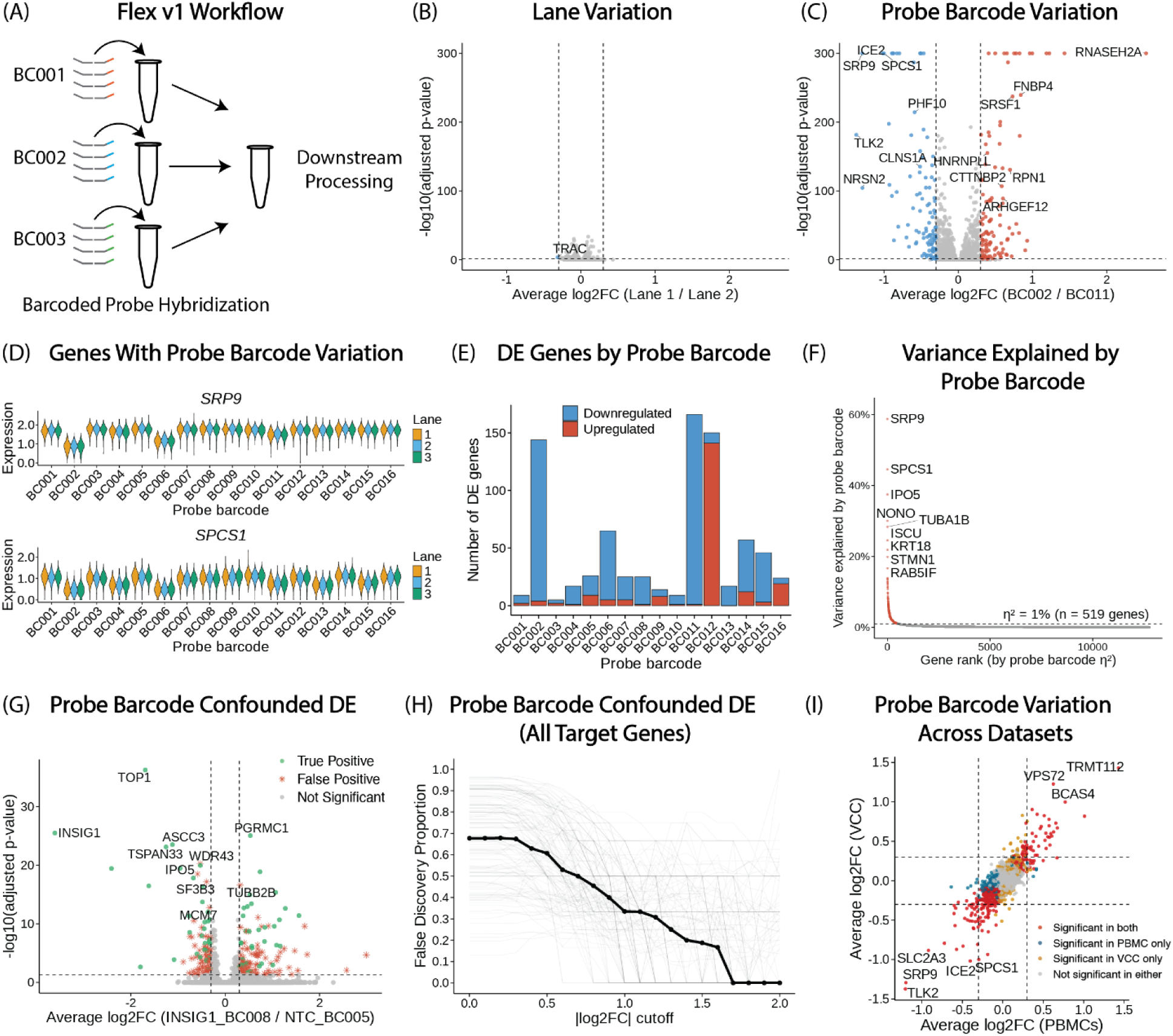
Probe barcode variation in Flex v1. (A) Schematic of Flex v1 workflow. (B) Volcano plot of differential gene expression analysis between lane 1 and lane 2, within BC001 in the Arc Virtual Cell Challenge dataset. (C) Volcano plot of differential gene expression analysis between BC002 and BC011, within lane 1. (D) Violin plots of *SRP9* and *SPCS1* expression. (E) Barplot of differential gene expression analysis between each probe set barcode and all other probe set barcodes (lane 1 cells only), where a gene is considered differentially expressed if |log2FC| > 0.3 and adjusted *p* < 0.05. (F) Gene rank plot of percentage of variance explained by probe set barcode. (G) Volcano plot of confounded differential gene expression analysis between INSIG1 guide cells and non-targeting control (NTC) guide cells, confounded by BC008 and BC005, which are two average effect barcodes. (H) Line plot of false discovery proportion across log2FC thresholds for BC008 and BC005 confounded analysis. Each light grey line represents a guide versus non-targeting control (n = 139 target genes). The dark black line is the median across all target genes. (I) Scatter plot showing BC002 versus BC011 differential gene expression analysis between the Arc Virtual Cell Challenge (VCC) dataset and the 10x Genomics PBMC dataset.

To test the assumption that probe set barcode variation is minimal in the Flex v1 assay, we analyzed a large scale Perturb-seq dataset generated by the Arc Institute for the 2025 Virtual Cell Challenge (VCC)^6^. In this experiment, the same population of cells subject to a pool of CRISPR perturbations was split randomly into 16 probe set barcode hybridization reactions, then loaded into three lanes for droplet generation and downstream profiling. Across all cells, in the absence of technical effects, there should be no differences in gene expression between lanes and between barcodes. This assumption holds true for lane-to-lane comparisons (Figure 1B), but not for comparisons across probe set barcodes (Figure 1C). Differential gene expression between probe set barcodes BC002 and BC011, within a single lane, reveals 217 differentially expressed genes (|log2FC| > 0.3, adjusted *p* < 0.05). This probe set barcode variation is strongly correlated between lanes within the dataset (Supplementary Figure 1A; Spearman ⍴ = 0.73, *p* < 0.01). Some genes are differentially expressed across multiple probe set barcodes, while others are differentially expressed in single probe set barcodes (Figure 1D). The probe set barcode variation is not explained by a single problematic barcode, as 5 of the 16 barcodes have more than 50 differentially expressed genes when compared to all other barcodes (Figure 1E; |log2FC| > 0.3, adjusted *p* < 0.05).

Assessing Flex v1 probe set barcode batch effect by UMAP visualization does not reveal any obvious differences in the projected embedding space (Supplementary Figure 1B). Standard quality control metrics are generally the same across probe set barcodes (Supplementary Figure 1C-E). However, we did note some probe set barcodes, such as BC002, with small but notable differences in mean UMI counts per cell. We wondered if these differences could be driving some of the probe set barcode variation. Controlling for the number of UMIs per cell in a logistic regression differential expression analysis between BC002 and BC011 did not correct the variation observed with the Wilcoxon Rank Sum test (Supplementary Figure 1F; Spearman ⍴ = 0.82, *p* < 0.01). To account for the method of differential expression analysis, we tested 7 common differential expression methods deployed by Seurat and observed very similar results between methods (Supplementary Figure 1G). Overall, for 519 genes, greater than 1% of variance in gene expression can be explained by probe set barcode when subsetting to non-targeting control (NTC) cells (Figure 1F). By comparison, lane identity explains greater than 1% of variance in gene expression for only 27 genes in the same NTC cells (Supplementary Figure 2A). Importantly, percent variance in gene expression explained by probe set barcode is positively correlated with mean gene expression (Supplementary Figure 2B). Altering the threshold for the minimum percentage of cells expressing a gene for it to be considered differentially expressed did not meaningfully change the probe set barcode variation observed (Supplementary Figure 2C).

The use of Flex (v1) probe set barcode as a biological sample barcode raised concern that technical batch effect could confound case-control comparisons, leading to the identification of false positive differentially expressed genes. Many cytokine perturbations produce response signatures of ∼10-100 differentially expressed genes when defining a differentially expressed gene as one with absolute log2FC greater than 0.25 and FDR-adjusted *p* less than 0.05 using the Wilcoxon Rank Sum test^8,9^. To test how different probe set barcodes in Flex v1 may confound case-control analyses, we leveraged the orthogonally barcoded guides in the VCC Perturb-seq dataset. We first established a ground truth of differentially expressed genes by comparing each guide (n = 150 target genes) to the NTC cells within a probe set barcode. We then deliberately confounded guide identity with probe set barcode by comparing NTC cells from one probe set barcode to target gene guide cells from a different probe set barcode. Because we had already established a ground truth set of differentially expressed genes for each guide (using within-barcode comparisons), any additional genes that appeared as significant in the confounded analysis could be attributed to probe set barcode batch effect rather than the guide perturbation itself. This design allowed us to directly quantify false positive rates introduced by probe set barcode confounding. Taking the probe set barcodes that produce an average technical effect amongst the 16 tested, we compared targeting cells from BC008 to nontargeting guide cells from BC005. For INSIG1 knock-out cells versus nontargeting cells, 172 genes were false positives, making up 71% of the differentially expressed genes (Figure 1G). The false positive genes were not obviously distinguishable from true positive genes by effect size or by *p* value.

Generalizing this analysis across all guides versus non-targeting controls using BC008 and BC005 as confounders, we find at an absolute log2FC threshold of 0.3 that a median of 67% of differentially expressed genes are false positives (Figure 1H). The percentage of genes that are false positive decreases to 34% at a log2FC threshold of 1, only falling to 0% at a log2FC threshold of 1.7. For guides that produce many true differentially expressed genes, the proportion of genes that are false positive is lower (Supplementary Figure 2D). For the worst case scenario probe set barcode pair in the dataset, the percentages of false positive genes are even more concerning. When introducing BC011 and BC002 as confounders, at a log2FC threshold of 0.3 a median of 91% of the differentially expressed genes are false positives. The percentage of genes that are false positives only decreases to 50% at a log2FC threshold of 2 (Supplementary Figure 2E-F). For the two probe set barcodes that produce the smallest technical effect, BC003 and BC010, the median false discovery proportion is 46% at a log2FC threshold of 0.3 and 20% at log2FC equal to 1 (Supplementary Figure 2G-H). Overall, if the Virtual Cell Challenge dataset barcoded perturbations with single probe set barcodes, many of the differentially expressed genes would have been false positives.

This probe set barcode variation effect is not unique to the Virtual Cell Challenge dataset. We also discovered significant variation across probe set barcodes in a 16-plex PBMC dataset published by 10x Genomics (Supplementary Figure 2I). The barcodes with the strongest batch effects vary between datasets, but BC002, BC006, and BC011 are consistently among the worst offenders. Interestingly, while there are some genes that seem context dependent in their probe set barcode variation, many of the genes that are differentially expressed between BC002 and BC011 in the Virtual Cell Challenge dataset are also differentially expressed between the same probe set barcodes in the PBMC data, suggesting the technical effect holds true across datasets and contexts (Figure 1I, Supplementary Figure 3). The presence of consistent probe set barcode effects across multiple datasets introduces uncertainty over whether probe set barcodes in Flex v1 can truly serve as sample barcodes. Without any further correction, such artifacts make differential expression comparisons across probe set barcodes challenging to interpret.

10x Genomics recently released Flex v2, an updated version of the probe-based technology. In contrast to Flex v1, where the probe barcode is embedded directly within the transcript probes and is inseparable from the hybridization reaction, Flex v2 decouples sample barcoding from probe hybridization entirely (Figure 2A). In Flex v2, all samples are hybridized with the same barcode-free whole transcriptome probe set. Sample identity is instead conferred by a separate barcoding oligo hybridization step performed after probe hybridization is complete.

**Figure 2.**
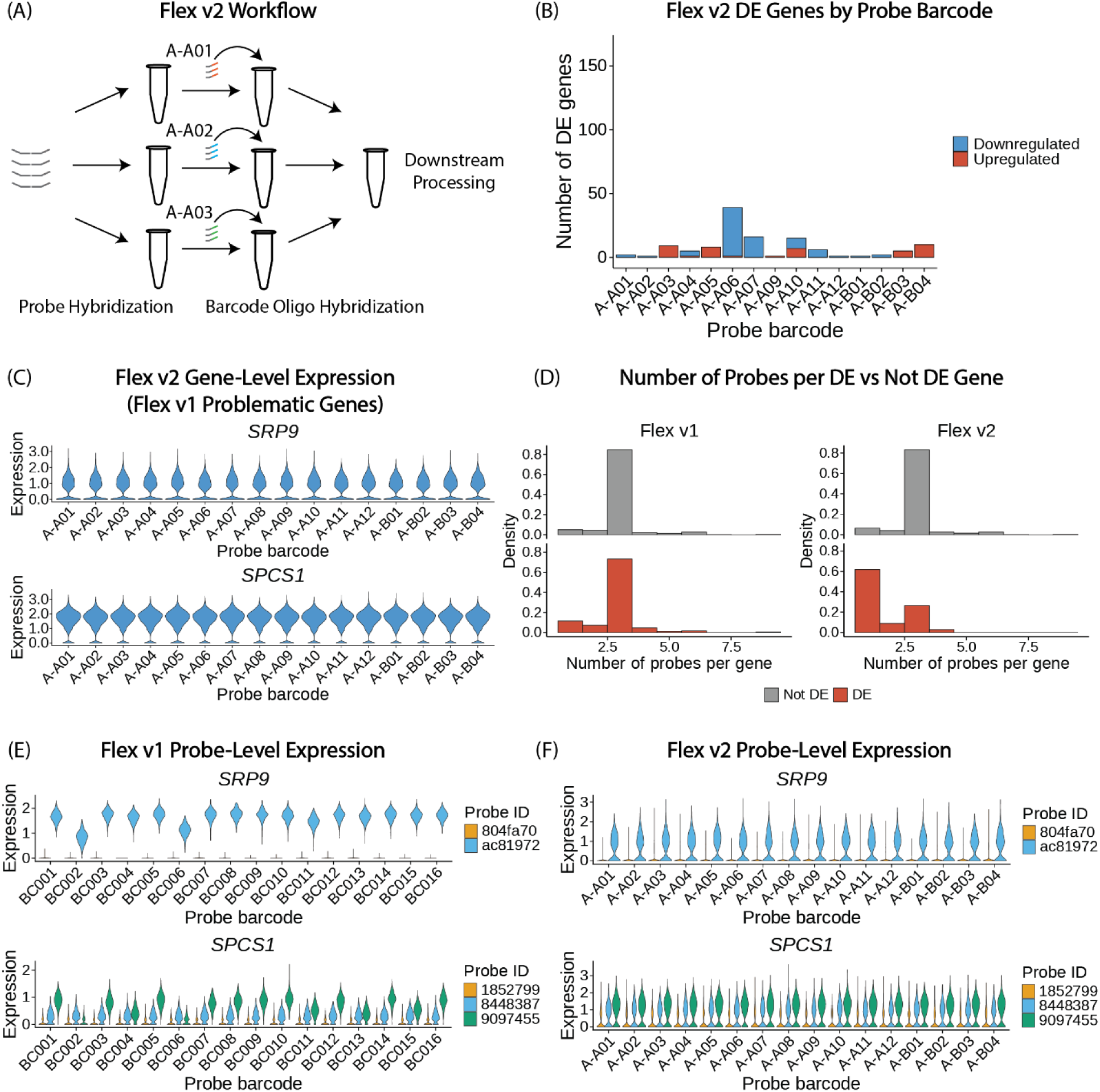
Probe barcode variation in Flex v2. (A) Schematic of Flex v2 workflow. (B) Barplot of differential gene expression analysis between each probe barcode and all other probe barcodes in the A375 Flex v2 dataset across all cells, where a gene is considered differentially expressed if |log2FC| > 0.3 and *p* < 0.05. The y axis is set to the same range as the y axis in Figure 1E for comparison with Flex v1. (C) Violin plots of *SRP9* and *SPCS1* expression. (D) Histograms of the distribution of probes per gene split by DE status for the Flex v1 VCC dataset (non-targeting control cells in lane 1) and for the Flex v2 A375 dataset (PBS-treated cells only). (E) Violin plots of *SRP9* and *SPCS1* expression split by individual probes for the Flex v1 VCC dataset (non-targeting control cells in lane 1). (F) Violin plots of *SRP9* and *SPCS1* expression split by individual probes for the Flex v2 A375 dataset (PBS-treated cells only).

We designed an experiment to test Flex v2 where we split a population of A375 cells evenly across 16 probe barcodes. Importantly, we split the population into 16 independent reactions for the probe hybridization step, which simulates a 16-sample experiment. Quality control metrics were comparable across probe barcodes (Supplementary Figure 4A-C). Probe barcode A-A06 had 39 differentially expressed genes when compared against all other probe barcodes, while all other tested probe barcodes had fewer than 20 differentially expressed genes (Figure 2B; |log2FC| > 0.3, *p* < 0.05). Of the 52 total differentially expressed genes in any probe barcode in Flex v2, 16 of them overlap with the 563 total differentially expressed genes by any probe barcode in the VCC Flex v1 dataset. The most concerning genes in Flex v1 did not vary by probe barcode in Flex v2 (Figure 2C). While we still note some differentially expressed genes across probe barcodes (Supplementary Figure 4D-E), the overall level of probe barcode variation is lower than we observed in Flex v1. Notably, the number of differentially expressed genes between probe barcodes in Flex v2 approaches zero when the initial probe hybridization is performed in one pooled reaction before the cells are split randomly for the barcoding oligo hybridization step (Supplementary Figure 5). This is promising for superloading one sample but does not represent a sample-barcoded Flex v2 experiment.

Flex v2 reduces the probe barcode effect observed in Flex v1 but the mechanism underlying these differences remains unclear. At the global probe barcode level, we noted small but significant differences in the number of UMIs sequenced per cell in some of the problematic probe barcodes. However, controlling for UMIs per cell in the differential gene expression analysis did not correct the effect. At the gene-level, the problematic genes were not obviously enriched for genes associated with cell quality, like mitochondrial genes. While there was some positive correlation between variance explained by probe barcode and mean expression, this was not a convincing explanation for the observed technical probe barcode effect.

For a better mechanistic explanation, we looked at the probe-level data to explore several hypotheses, including probe set design, probe-to-probe barcode secondary structure, and probe-to-probe correlation within a gene. First, the probe sets are nearly identical between v1 and v2, so the probe sequences themselves do not explain the difference. Second, since the Flex v1 assay performs the probe hybridization step with pre-barcoded probes, while Flex v2 decouples the probe hybridization from the probe barcoding, we wondered if the probe barcode could be physically interacting with the probe hybridization region to interfere with transcript binding. While each of the 16 probe barcodes in Flex v1 are composed of four unique 8 bp sequences as well as 1 bp frameshift sequences to account for sequencing errors, in Flex v2, there is only one unique 10 bp sequence for each of the 384 probe barcodes. However, while there were differences in UMI counts between different probe barcode sequences for individual probe barcodes, the overall pattern of variable expression for these problematic genes in particular barcodes is retained across probe barcode sequences (Supplementary Figure 6A-B). Therefore, given that the expression patterns are broadly similar across probe barcode sequences, this explanation is unlikely.

Third, we found that in both Flex v1 and v2, differentially expressed genes between probe barcodes had different numbers of probes per gene, with single-probe genes enriched in the set of problematic genes (Figure 2D). This aligns well with our previous observation that differentially expressed genes between probe barcodes tend to be highly expressed as Flex addresses the read disparity between highly and lowly expressed genes by only designing one probe pair for highly expressed and mitochondrial genes (Supplementary Figure 2B). For the problematic genes with multiple probes, we considered whether their variance could be explained by variable expression between probes. In both Flex v1 and v2, we found that generally, across all genes, there tended to be many genes where at least one probe was expressed in fewer than 10 percent of cells, a phenomenon we termed probe dropout (Supplementary Figure 7A-B). This meant that despite genes being targeted by more than one unique probe sequence, at least one of those probes did not recover any gene expression. Therefore, sometimes a gene might be targeted by more than one probe, but in reality be more similar to a single-probe gene. We found that in both Flex v1 and Flex v2, differentially expressed genes tended to be more likely to have at least one dropout probe, although this effect was not significant (Supplementary Figure 7C-D). This led us to wonder whether expression changes in individual probes between probe barcodes were responsible for the gene-level effect or whether all of a gene’s probes exhibited similar expression changes across probe barcodes. Within problematic genes, we found that expression changes in individual probes were often responsible for the gene-level effect (Figure 2E-F, Supplementary Figure 7E-G). We quantified this by comparing the differentially expressed probes and genes across probe barcodes identified using the Wilcoxon Rank Sum test (Supplementary Figure 7E-F). The vast majority of the differentially expressed multi-probe genes had one differentially expressed probe. In rare cases, more than one probe exhibited expression differences across probe barcodes (Supplementary Figure 7E-G). These results indicate that probe-level differences are responsible for the technical variation across probe barcodes in Flex v1. Our previous analysis rules out several explanations tied to the unique characteristics of the probe barcode and probe sequence themselves. Therefore, we believe batch effects in the manufacturing of separate barcoded probe sets, such as differences in probe concentrations and probe truncations, could account for much of the observed differences.

In summary, we demonstrate that probe set barcodes are a significant source of technical variation in the Flex v1 assay. In a dataset where the same population of cells was split across 16 probe set barcode hybridization reactions, hundreds of genes are differentially expressed between probe set barcodes, with the effect reproducible across lanes and consistent across independent datasets. When probe set barcode identity is confounded with biological sample identity, a large proportion of differentially expressed genes at standard thresholds are false positive genes. The problem is not attributable to a single outlier barcode, is not corrected by controlling for UMI depth, and is robust across differential expression methods. While increasing the log2FC threshold does decrease the proportion of false positives, many single cell RNA sequencing analyses rely on moderate-to-low effect size differences in gene expression to derive robust transcriptional programs and perform gene set enrichment analysis. The Flex v2 assay, which decouples sample barcoding from probe hybridization, eliminates probe set barcode manufacturing as a possible batch effect. While we still observe some small technical variation between independent probe hybridization reactions, we attribute this to experimental variability in sample processing since each hybridization reaction uses probes from the same probe set. Overall, the sample-to-sample technical variation is significantly reduced compared with Flex v1.

These findings carry direct implications for experimental design. Users of Flex should treat probe barcodes as potential confounding variables and be cautious when using it as a biological sample barcode for case-control comparisons. We conclude that Flex v2 is much more suitable than Flex v1 for designs that rely on probe barcodes as a biological sample barcode. Replicates are recommended to mitigate probe barcode technical effects. We did not assess mouse probe sets in this study but recommend the same level of scrutiny for Flex v1. For human datasets that have already been generated with Flex v1, we encourage examination of results for the specific genes we flagged as problematic in this study before drawing biological conclusions that rely on comparisons across probe barcodes (Supplementary Table 1-3).

The significance of these observations is extended by the rapid adoption of probe-based assays for atlas-scale data collection and virtual cell model training. Datasets that do not compare directly across probe set barcodes, like those generated by many Perturb-seq style projects, should be largely unaffected by these artifacts. More broadly, these findings offer a lesson in the development of probe-based single cell technologies and the analysis of single cell data. The probe set barcode batch effects observed here in Flex v1 are substantially stronger than the lane-to-lane variation associated with capture-based single-cell technologies, especially when the data was collected in the same batch on the same microfluidic chip. These technical effects seem to be greatly reduced in Flex v2. As more probe-based data is generated and analysed, probe hybridization and sample barcoding should be considered as sources of technical variation during experimental design and analysis. Further, as future probe-based assays are developed, these technical effects should be considered in the chemistry and workflow of the technology to ensure the scalability and sensitivity potential of probe-based single cell profiling is translated into meaningful biological discoveries.

## Methods

### Preprocessing and normalization

All of the gene level analyses on the Virtual Cell Challenge dataset were performed on the training dataset published for the competition. For probe-level analyses, *cellranger multi* (v.9.0.1) was run on fastq files from lane 1 of the experiment to generate the molecule_info.h5 file. For Flex v2, we used *cellranger multi* (v.10.0.0) to process the data with default parameters. The Cell Ranger results were then loaded into R (v.4.5.2) using Seurat (v.5.4.0). For the Flex v1 PBMC dataset and the Flex v2 A375 dataset, cells were filtered to those with at least 200 genes and below 5% mitochondrial reads. Data was normalized and log-transformed with a global-scaling normalization method with *NormalizeData* in Seurat using the default values (‘LogNormalize’ and a scale factor of 10,000). For the probe-level analysis, the raw_probe_bc_matrix output from Cell Ranger was filtered to the same cell barcodes that remained following filtered_feature_bc_matrix preprocessing and was normalized as described above.

### Differential gene expression

To identify differentially expressed genes across all probe barcodes, Seurat’s *FindAllMarkers* was performed using the Wilcoxon method and a min.pct of 0.1 and logfc.threshold of 0, unless otherwise specified. For the comparison of differentially expressed genes across different methods, the min.pct remained 0.1 with all other parameters set to their default. For individual probe-barcode-to-probe-barcode differential expression, Seurat’s *FindMarkers* was used. We also tested using *scanpy* (v.1.11.5) and Python (v.3.11.14) for differential expression and observed similar results (data not shown). We used the same approach on the probe level matrices to identify differentially expressed probes across all probe barcodes.

### Variance explained analysis

To quantify the fraction of gene expression variance attributable to probe barcode identity, we computed eta-squared (η² = SS_between / SS_total) from a one-way ANOVA for each gene, using probe barcode (K = 16) as the grouping factor. This analysis was restricted to NTC cells (∼38,000 cells across 3 sequencing lanes). Only genes detected in ≥5% of NTC cells were tested, and the same calculation was repeated with sequencing lane (K = 3) as the grouping factor for comparison. Effect size thresholds (η² ≥ 1%) were used as the primary interpretive metric.

### Probe barcode confounded differential expression analysis

To quantify the false positive rate introduced by probe barcode confounding in case-control comparisons, we leveraged the orthogonally barcoded guide library in the Virtual Cell Challenge Perturb-seq dataset, which contains cells from 150 target genes and NTC guides distributed across 16 probe barcodes. For each of the 150 target genes, a ground truth set of differentially expressed genes was defined by comparing cells carrying that guide to NTC cells within the same probe barcode (within-barcode comparison). Differential expression was performed using the Wilcoxon rank-sum test (Seurat *FindMarkers*, test.use = "wilcox", min.pct = 0.1, logfc.threshold = 0.0), and genes were considered significant at FDR-adjusted *p* < 0.05 and absolute log2FC greater than or equal to a given threshold. Guides with fewer than 20 cells in the focal probe barcode were excluded from analysis. To simulate confounding of guide identity with probe barcode, a second DE analysis was performed by comparing guide-bearing cells from one probe barcode to NTC cells from a different probe barcode (cross-barcode comparison) using identical test parameters. The cross-barcode comparison was performed for three barcode pairs selected to represent worst-case (BC002 vs. BC011), average (BC005 vs. BC008), and smallest (BC003 vs. BC010) pairwise technical effects among the 16 probe barcodes, based on the magnitude of between-barcode expression differences established in preceding analyses. For each guide, the false discovery proportion at a given log2FC threshold was defined as the number of genes significant in the cross-barcode comparison but absent from the ground truth set, divided by the total number of genes significant in the cross-barcode comparison. Genes significant in the confounded analysis but absent from the ground truth were classified as false positives attributable to probe barcode batch effect. The false discovery proportion was calculated across a sweep of absolute log2FC thresholds (0.1 to 2.0 in increments of 0.1) to characterize how stringent effect-size filtering affects the proportion of false positives, and median false discovery proportion across all guides is reported at each threshold.

### Distribution of number of probes per gene analysis

The number of probes ids per gene were determined using the v2.0 and v1.1.0 probe sets provided by 10x Genomics and the distribution of number of probe ids per gene observed for genes output as differentially expressed or not differentially expressed from the differential gene expression analysis described above for the lane 1 non-targeting control cells in the Flex v1 VCC dataset and the PBS treated cells in the Flex v2 dataset.

### Probe barcode sequence variation analysis

Reads were extracted from the Cell Ranger-aligned Binary Alignment Map (BAM) file genes of interest, including SRP9 and SPCS1 using samtools (1.21) view filtered by the gene name (GN) tag and restricted to the gene’s genomic coordinates. For each read, the following fields were parsed from the SAM optional tags: cell barcode (CB, 24 bp), uncorrected probe barcode (last 8 bp of CR tag), corrected probe barcode (last 8 bp of CB tag), probe ID (pr tag), and UMI (UB tag). Reads were filtered to retain only those originating from cell barcodes of non-targeting control cells in lane 1 of the VCC dataset. Multi-probe read annotations containing ";NA" or "NA;" in the pr tag were cleaned by removing the NA portion. Each uncorrected 8 bp probe barcode sequence was mapped to the probe barcode ID and the sequence version identified using the mapping list provided by 10x Genomics: probe-barcodes-fixed-rna-profiling-rna.txt. UMIs were deduplicated per unique combination of cell barcode, UMI sequence, and probe ID. Per-cell UMI counts were then aggregated by cell barcode, gene, probe ID, raw probe barcode, and corrected probe barcode.

### Probe Dropout Analysis

To assess probe-level variability in the Flex assay, we analyzed individual probe detection rates across cells. For each probe in the set, we computed the fraction of cells in which the probe was detected (count > 0) and its mean expression level from the probe-level Seurat object. Probes were classified as "dropout" if they were detected in fewer than 10% of cells. For genes targeted by two or more probes, we summarized the number of dropout probes per gene and classified each gene by its dropout count. Differentially expressed genes were designated as described previously

### Flex v2 A375 dataset generation

Eight wells of 150,000 A375 cells were treated with either one of seven ligands (IFNG, TNFA, GDF15, SPP1, POSTN, ANXA1, and MDK) or PBS as a control for 24 hours. Cells were then washed with PBS, trypsinized, centrifuged at 400 x g for three minutes, and resuspended in 1% BSA in PBS. Following another centrifugation, cells were resuspended in 45 uL of 1% BSA in PBS with 5 uL of Human TruStain FcX and incubated for 10 minutes at 4℃. After adding another 50 uL of 1% BSA in PBS, 0.1 ug of TotalSeq™-C antibody was added to each tube. Following another 30 minute incubation on ice, the cells were washed according to 10x’s CG000781 protocol. Although cells from each sample were counted, samples were pooled equally into a Protein LoBind Tube. Samples were fixed according to 10x’s CG000782 protocol. The rest of the sample preparation for sequencing followed 10x’s CG000835 protocol (Rev B) for GEM-X Flex v2 with Feature Barcode technology for Protein. The 1.5 million cells counted after fixation were split evenly into 16 tubes for probe hybridization. Following the barcode oligo hybridization, samples were washed in pooled format. The entirety of each sample was added to the pool. Of the 1.4 million cells recovered following post-hybridization washing and resuspension, 232,000 cells were loaded for GEM generation and sequenced on a NovaSeqX 25B (11,897 reads per cell). Apart from Figure 2b, which was performed on all sequenced cells from the dataset, all other Flex v2 A375 analyses were performed on PBS-treated cells only.

### Flex v2 PBMC dataset generation

10x Genomics provided us with the *cellranger multi* filtered_feature_matrix output for three independent lots of the Set A barcode oligo plates tested on human PBMCs. The entire sample was hybridized to the whole transcriptome probe set in the same reaction before being split into 96 different barcode oligo hybridization reactions. Preprocessing, normalization, and differential gene expression analysis was performed as described previously.

## Data availability

We have deposited the A375 Flex v2 dataset on Zenodo along with probe-level data for the Virtual Cell Challenge dataset and the A375 dataset (https://zenodo.org/records/19363777). The Flex v1 Arc Institute Virtual Cell Challenge dataset is publicly available and we used the adata_Training.h5ad file (https://github.com/ArcInstitute/arc-virtual-cell-atlas). The Flex v1 PBMC dataset is publicly available and all relevant files can be downloaded directly from the 10x Genomics website (https://www.10xgenomics.com/datasets/320k_Human_PBMCs_Sub_Pool_16-plex_GEM-X_FLEX). Supplementary tables are available on GitHub (https://github.com/jacksonweir/flex-technical-variation).

## Code availability

The code to reproduce the main analyses and figures in this manuscript is available on GitHub (https://github.com/jacksonweir/flex-technical-variation). We also included a structured skill describing how AI coding agents should think about the source of technical variation explored in this paper with the goal of enabling readers to download the skill and quickly check their own Flex data for probe barcode-associated technical variation.

## Acknowledgements

We thank 10x Genomics for their support and assistance in designing and evaluating Flex v1 and Flex v2 experiments and providing Flex v2 data; P. Hsu for providing access to the raw data from the Virtual Cell Challenge dataset; C. Lareau, N. Hacohen, B. Cooper, M. Sade-Feldman, C. Chu, D. Lesmen, and the members of the Chen Laboratory for helpful discussions and feedback.

## Supplementary Figures

**Supplementary Figure 1.**
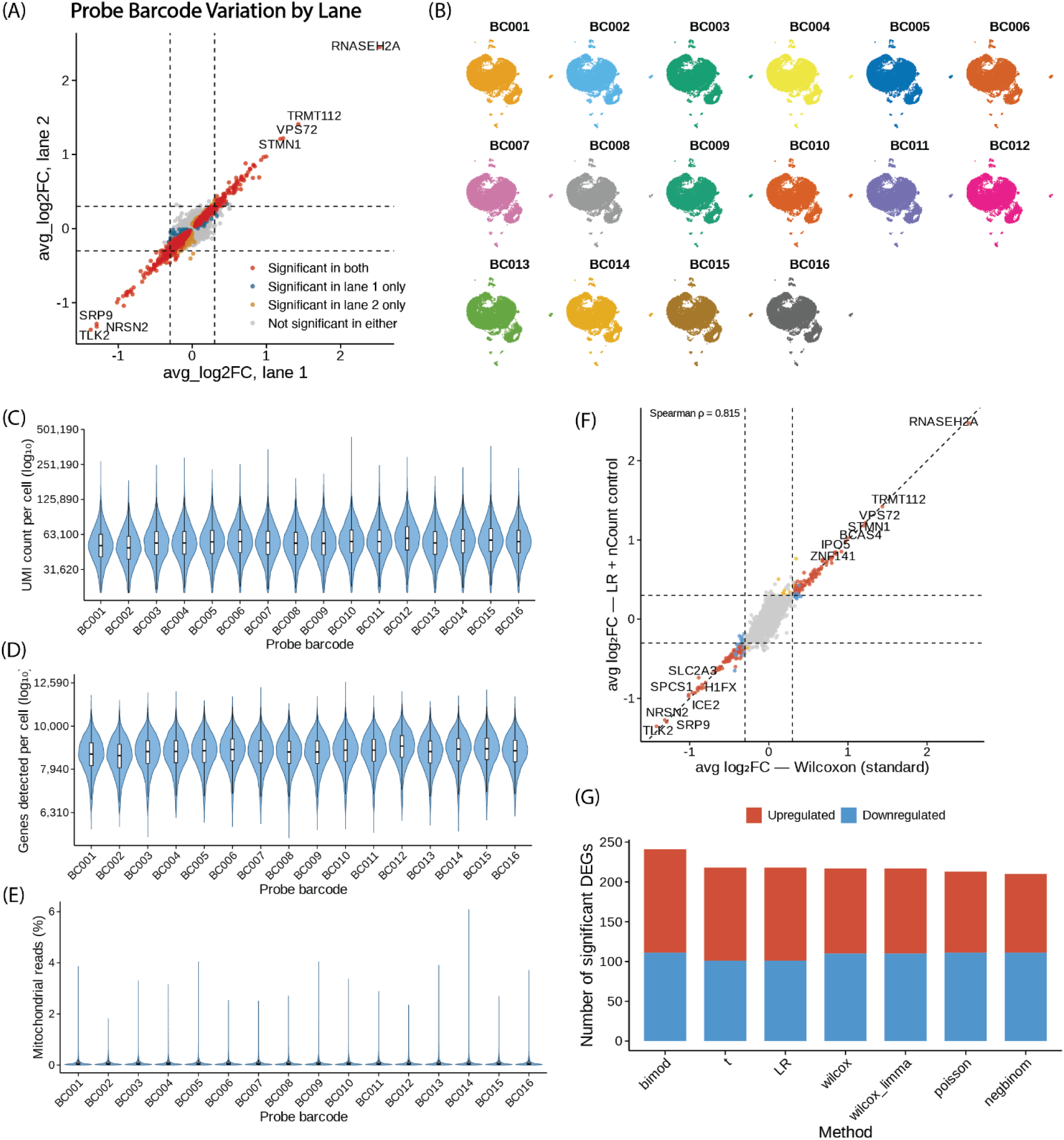
(A) Correlation between BC002 versus BC011 differential gene expression results between lane 1 and lane 2. (B) UMAP projection of profiled cells in the Arc Virtual Cell Challenge dataset split by probe set barcode. Violin plots split by probe set barcode for (C) UMIs per cell, (D) genes per cell, (E) mitochondrial reads per cell (%). (F) Scatter plot comparing BC002 vs BC011 differential gene expression results using the Wilcox Rank Sum test and a logistic regression controlling for UMIs per cell. (G) Barplot showing number of differentially expressed genes between BC002 and BC011 using different differential gene expression methods, where a significant gene has |log2FC| > 0.3 and adjusted *p* < 0.05.

**Supplementary Figure 2.**
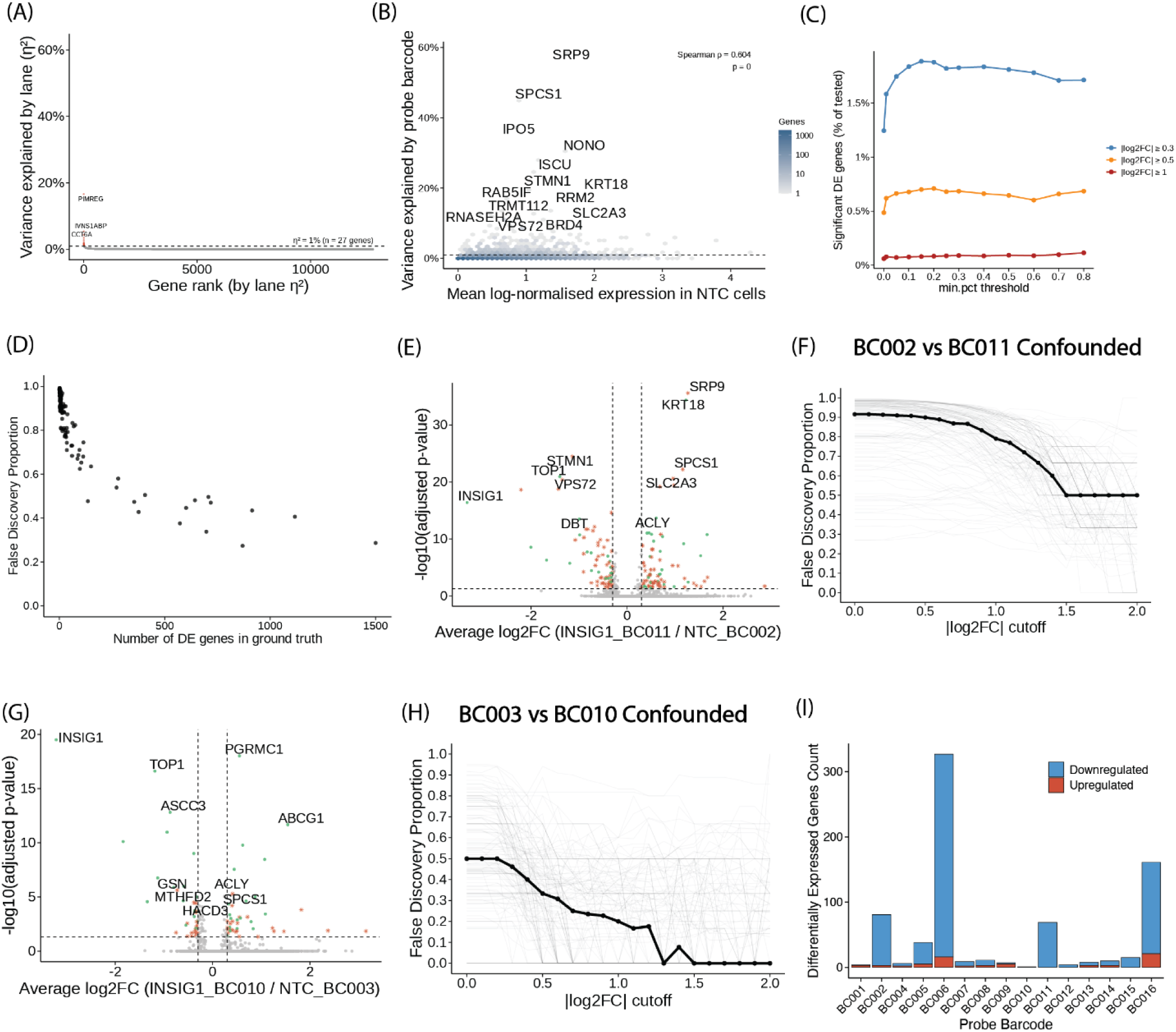
(A) Gene rank plot of variance explained by lane in the Virtual Cell Challenge dataset, with y-axis range matching Figure 1F. (B) Density scatter plot of variance explained by probe set barcode versus mean expression in non-targeting control cells. (C) Line plot showing the percentage of significant genes across |log2FC| thresholds and minimum cell expression thresholds (adjusted p < 0.05). (D) Scatter plot of false discovery proportion by number of ground-truth differentially expressed genes for BC011 and BC002 confounded analysis. (E) Volcano plot of confounded differential expression between INSIG1 and non-targeting control (NTC) guide cells, confounded by BC011 and BC002. (F) Line plot of false discovery proportion across log2FC thresholds for BC011 versus BC002. Volcano plot (G) and line plot (H) for BC003 versus BC010 confounded analysis. (I) Barplot of differential gene expression between each probe set barcode and all others (|log2FC| > 0.3, p < 0.05) in the 10x Genomics public 16-plex PBMC dataset.

**Supplementary Figure 3.**
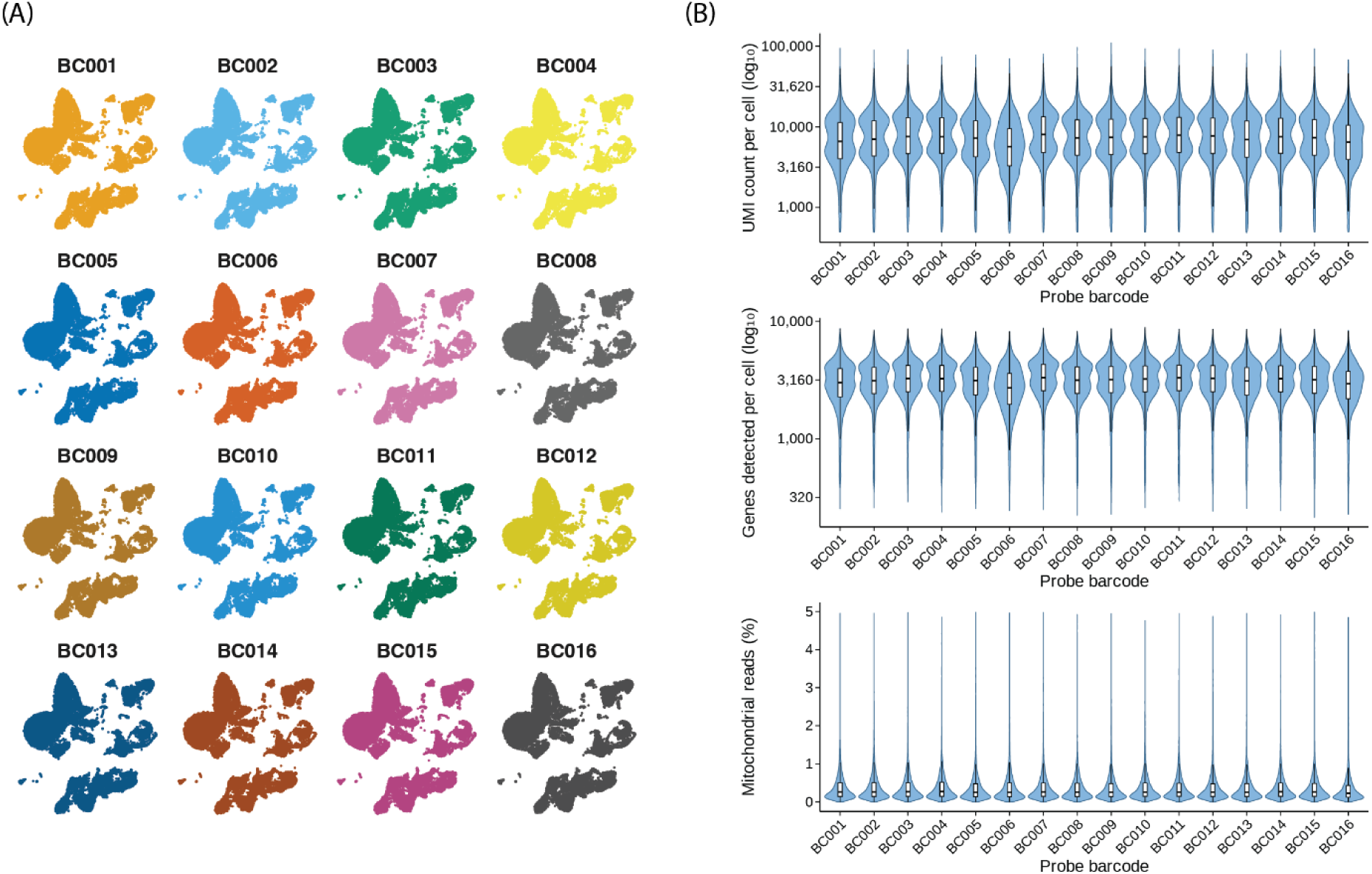
(A) UMAP projection of cells in the 10x Flex v1 PBMC dataset split by probe set barcode. (B) Violin plots split by probe set barcode for UMIs per cell, genes per cell, and mitochondrial reads per cell (%).

**Supplementary Figure 4.**
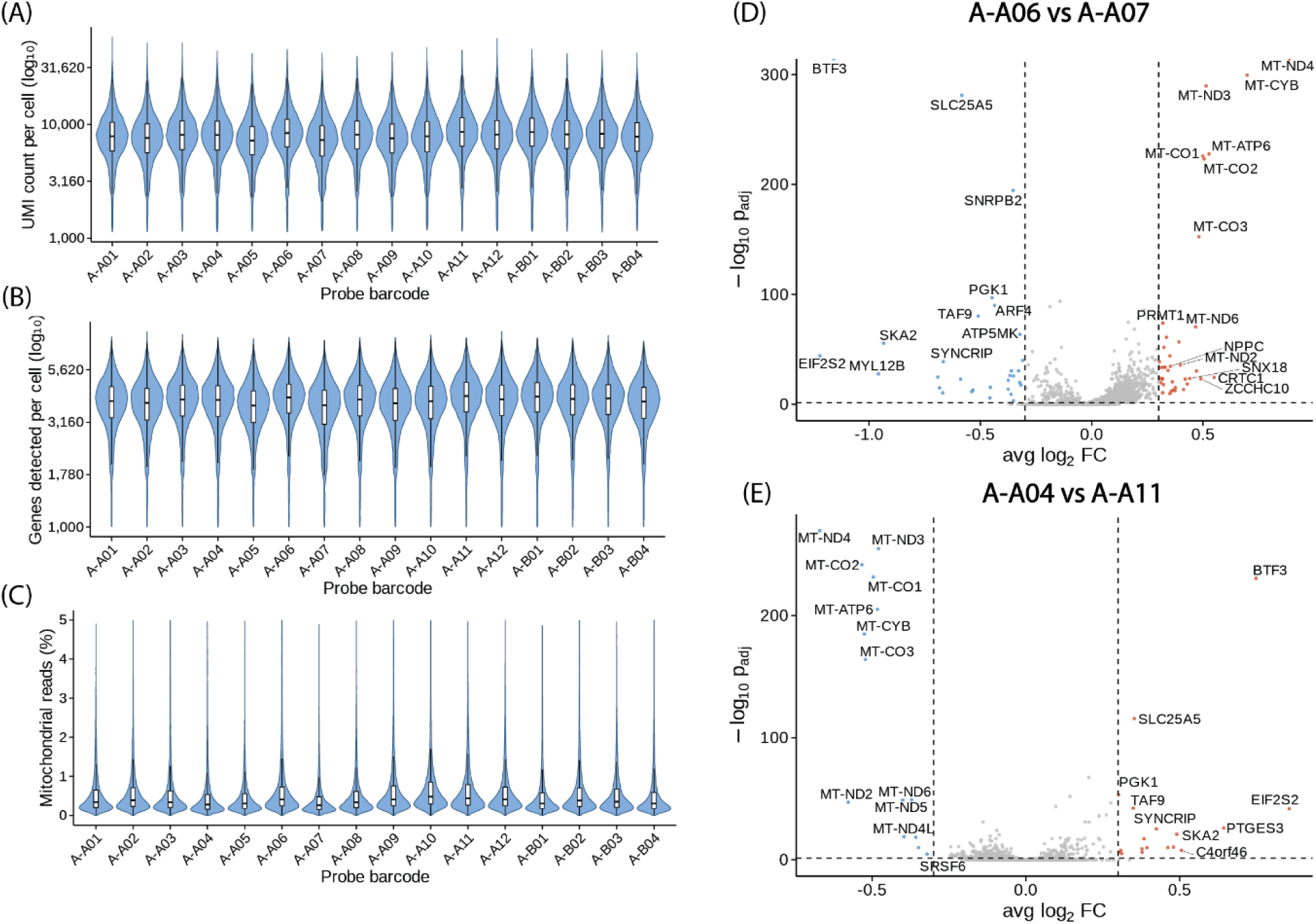
Violin plots split by probe barcode for (A) UMIs per cell, (B) genes per cell, (C) mitochondrial reads per cell (%) from the A375 Flex v2 dataset. Volcano plots of differential gene expression between (D) A-A06 and A-A07, and (E) A-A04 and A-A11.

**Supplementary Figure 5.**
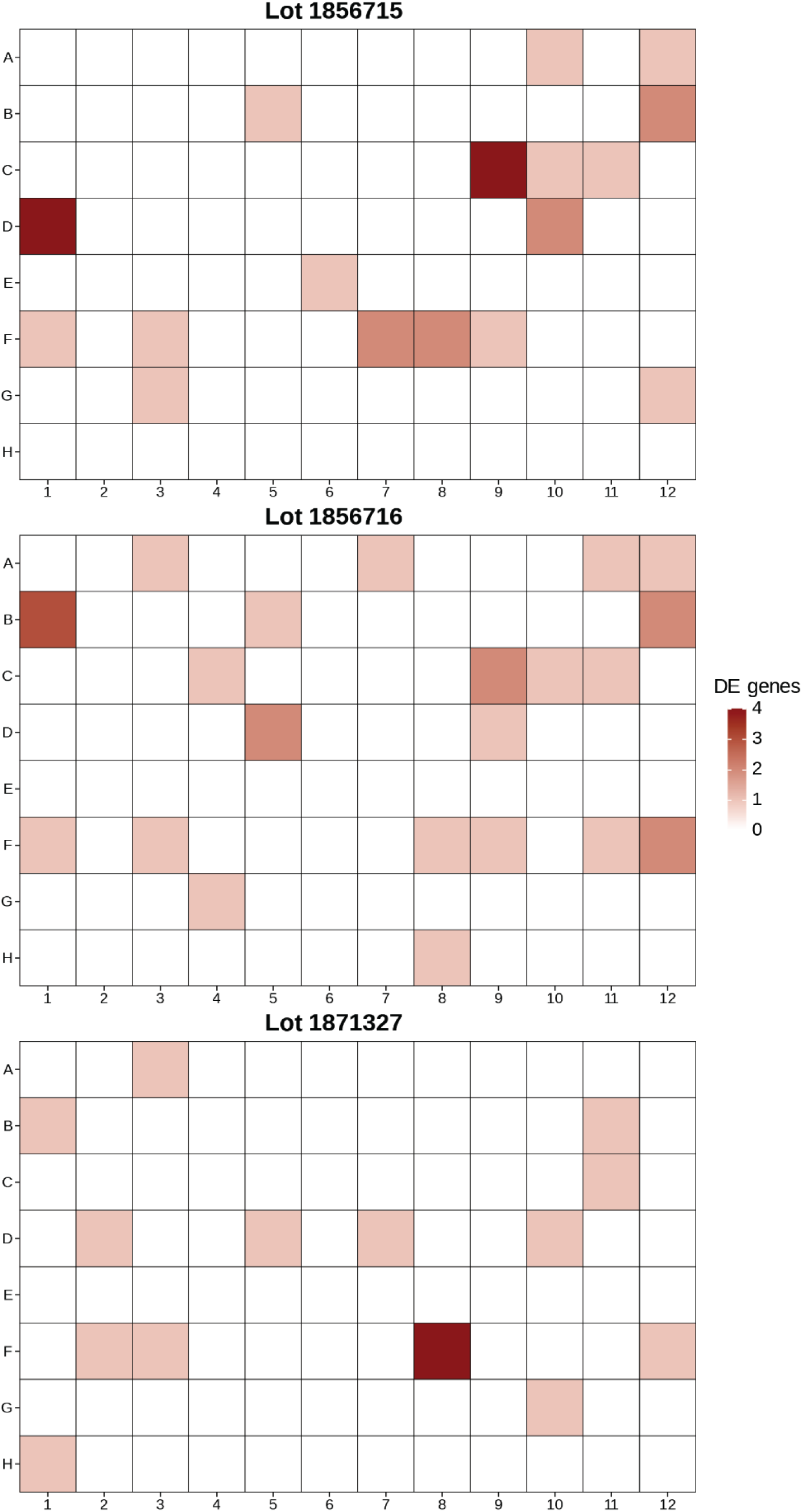
Human PBMC Flex v2 probe barcode differential gene expression results for three independent lots of the Set A barcode oligo plates. Differential gene expression was performed for cells of each probe barcode vs cells of all other probe barcodes within a lot. Unlike in the A375 Flex v2 dataset where probe hybridization was performed in separate tubes for each sample (each probe barcode), the probe hybridization was done in a single tube for this PBMC experiment before cells were split randomly into probe barcodes.

**Supplementary Figure 6.**
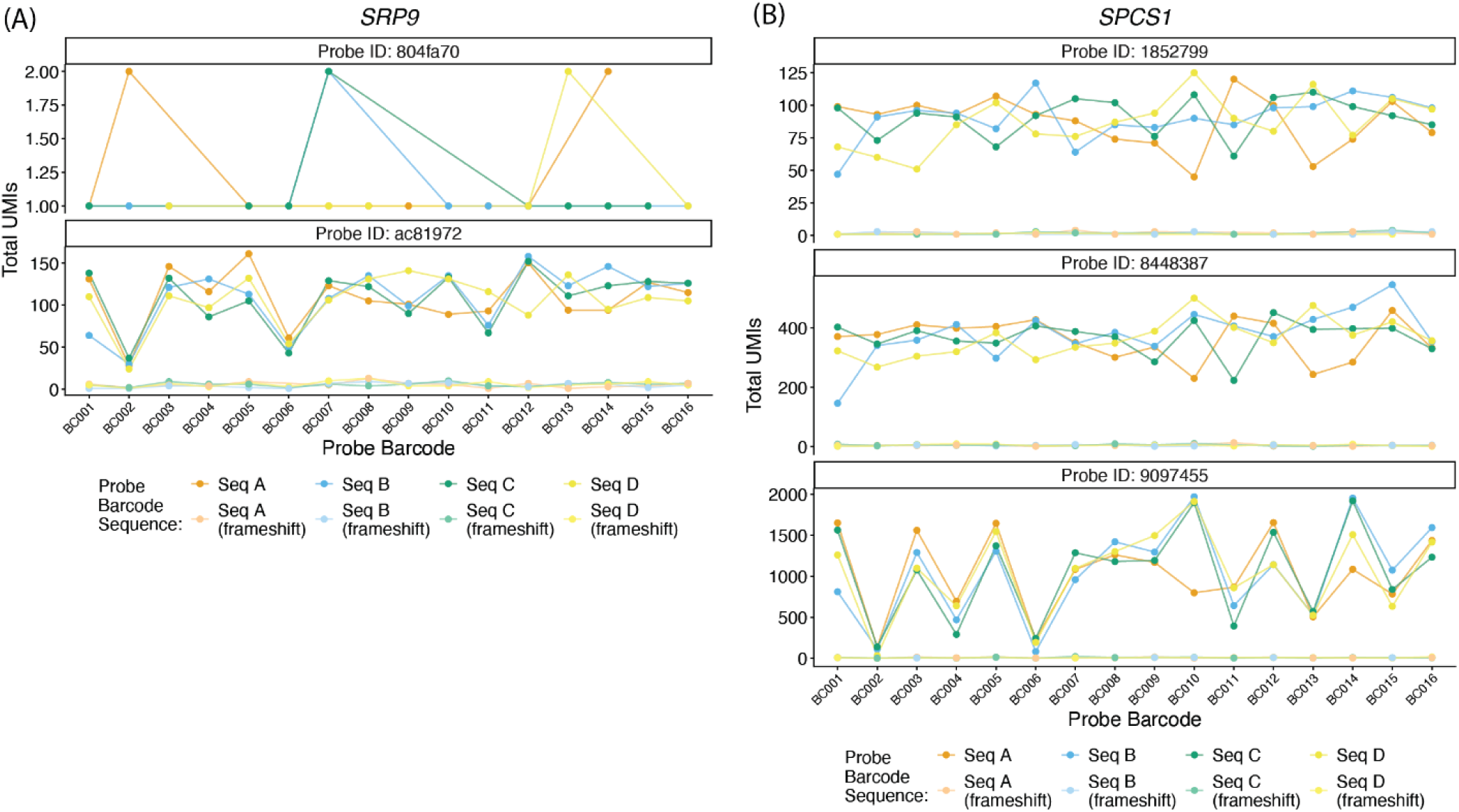
(A) Total UMI counts for each of the four probe set barcode sequence variants as well as their frameshift corrected sequence for each *SRP9* probe in the VCC dataset (lane 1, NTC cells only). (B) Total UMI counts for each of the four probe set barcode sequence variants as well as their frameshift corrected sequence for each *SPCS1* probe in the VCC dataset.

**Supplementary Figure 7.**
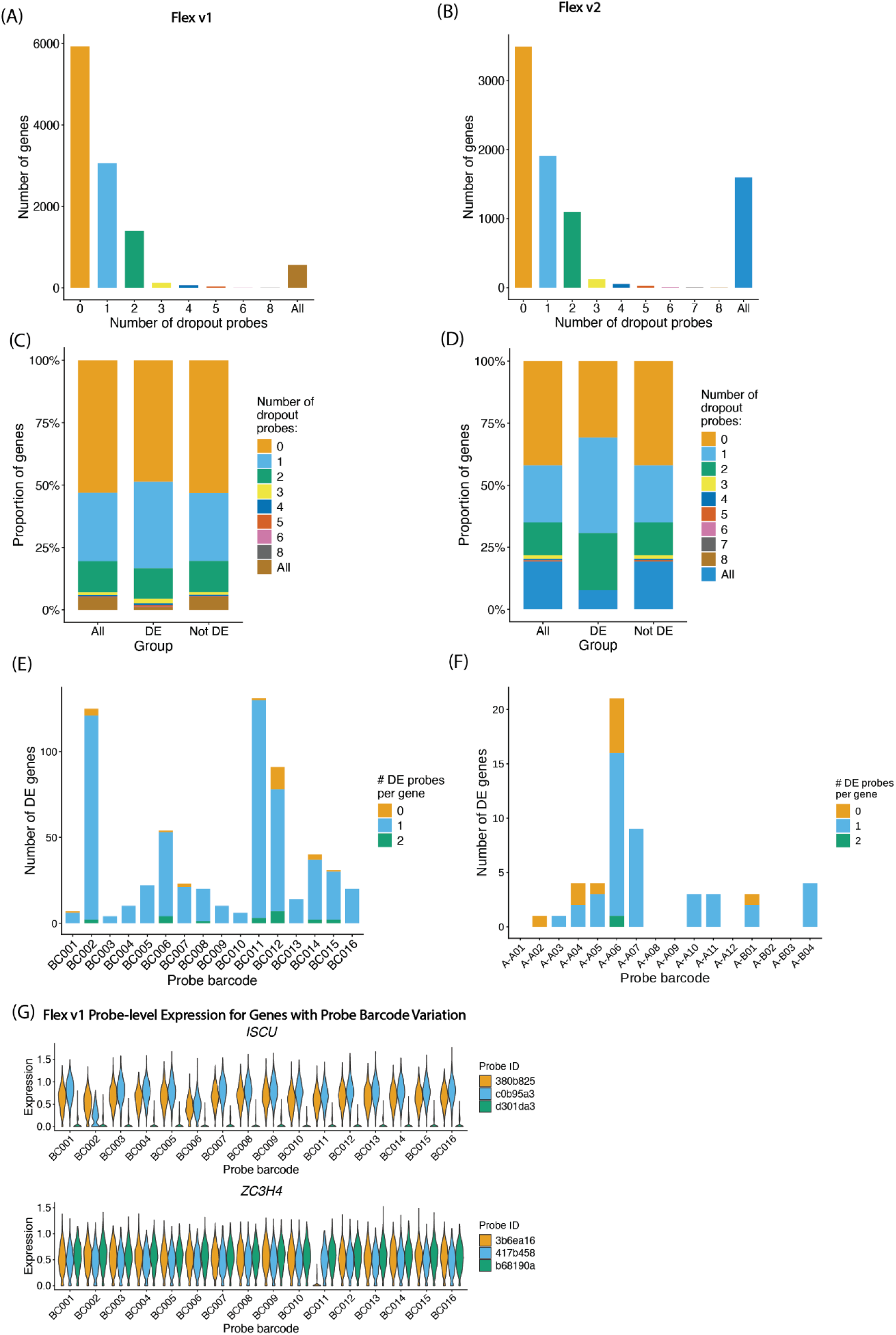
(A) Barplot of the number of probes expressed in less than 10 percent of cells (dropout probes) for all expressed genes in the Flex v1 VCC dataset (lane 1, NTC cells only). Expressed genes were identified as those expressed in greater than 10 percent of all cells. (B) Barplot of the number of dropout probes for all expressed genes in the Flex v2 A375 dataset (PBS-treated cells only). (C) Comparison of the proportion of genes with different numbers of dropout probes between all genes, differentially expressed genes, and not differentially expressed genes in the Flex v1 VCC dataset. Differentially expressed genes were identified as described previously. (D) Comparison of the proportion of genes with different numbers of dropout probes between all genes, differentially expressed genes, and not differentially expressed genes in the Flex v2 A375 dataset. (E) Number of DE probes for DE genes within each probe set barcode for the Flex v1 VCC dataset. (F) Number of DE probes for DE genes within each probe barcode for the Flex v2 A375 dataset. (G) Violin plots of *ISCU* and *ZC3H4* expression split by individual probes for the Flex v1 VCC dataset.

## Supplementary Tables

**Supplementary Table 1.** Results of one-vs-all differential expression analysis across probe set barcodes in the Virtual Cell Challenge (VCC) Flex v1 dataset (lane 1). Differential expression was performed using the Wilcoxon rank-sum test implemented in Seurat. Each row represents a gene tested in one probe set barcode group versus all others.

**Supplementary Table 2.** Results of one-vs-all differential expression analysis across probe set barcodes in a human PBMC Flex v1 dataset (all cells). Differential expression was performed using the Wilcoxon rank-sum test implemented in Seurat. Each row represents a gene tested in one probe set barcode group versus all others.

**Supplementary Table 3.** Results of one-vs-all differential expression analysis across probe barcodes in A375 melanoma cells processed with Flex v2 (all cells). Differential expression was performed using the Wilcoxon rank-sum test implemented in Seurat. Each row represents a gene tested in one probe barcode group versus all others.

